# Croaking for haste: How long does it take to describe a frog species since its discovery?

**DOI:** 10.1101/2025.04.17.649359

**Authors:** Albert Carné, Alberto Sánchez-Vialas, Claudia Lansac, Miriam Moreno, Ignacio De la Riva

**Author notes:** **Corresponding authors:** (AC), (IDLR).

## Abstract

Global biodiversity faces severe anthropogenic threats, with alarming extinction rates projected for the near future. Most of Earth’s diversity remains undescribed, meaning countless species are doomed to extinction before being documented. Since current conservation laws consider only described species, the time to achieve a representative inventory of global biodiversity is crucial for effective conservation. Amphibians, the most endangered vertebrate class, exemplify the challenge: while the number of threatened species rises, new species descriptions rapidly increase, and hundreds of candidate species are flagged annually worldwide. We analyzed all anuran species described from the year 2000 to 2023 across four biodiversity-rich tropical regions to investigate the time required to describe new frog species. We quantified the time needed to collect the type series, the number and timing of expeditions, the lag between collection and publication of the species description, and the total time. Additionally, we explored temporal trends and the effect of selected abiotic variables. On average, it takes 11.3 years to formally describe a frog species since the collection of the first specimen, with 4.5 years spent on specimen collection and 6.8 years on description and publication. These figures were consistent across three of the four regions analyzed. Alarmingly, the time required to describe new species is globally increasing, exacerbating the so-called taxonomic impediment. Only 36% of species were described within five years of collection, highlighting the importance of biological collections as reservoirs of undescribed diversity while also calling for specimen revision after expeditions. These results raise concerns about the effectiveness of current taxonomic and conservation practices in addressing the biodiversity crisis. We call for a global effort to prioritize taxonomic research and discuss innovative taxonomic and conservation approaches. Under current practices, and given the observed timelines, we will lose the race against extinction for many species.

## Introduction

Global biodiversity is being assaulted by Global Change. Populations are being reduced, contracted, and extirpated; species are becoming extinct at an unprecedented rate, and entire ecosystems are disappearing before we have the opportunity to properly prospect them [1–4]. The loss of a species is irreversible and its consequences are unpredictable [5]. The highest rates of decline and extinction are occurring in the tropics, hyper diverse areas where scientific knowledge remains limited, as they are known to contain high levels of undiscovered and undescribed diversity [3, 6–10].

Species are the fundamental unit to measure biodiversity. Only by recognizing the actual species richness of a given region can we discover large-scale ecological and evolutionary trends (e.g., [11]). How many species are on Earth remain an unresolved question — the Linnean shortfall [12–13], partly because of incomplete sampling and the lack of robust biodiversity characterization and extrapolation approaches [14–15]. However, there are three aspects of consensus: first, the largest proportion of biodiversity is comprised by undescribed species [15]. Second, the urgency to discover and describe this diversity has never been greater; naming species is the first step towards their conservation, and overcoming the Linnean shortfall is crucial for understanding distributional patterns and, consequently, for reaching effective conservation planning [16–17]. Third, we must explore new taxonomic approaches, as the estimated time required to document the remaining biodiversity far exceeds what may be available due to ongoing biodiversity threats [18–19].

Traditional taxonomic approaches, largely based on morphology, face significant challenges in addressing the urgency of the current biodiversity crisis. Integrative taxonomy emerged to accommodate and unify new concepts and methods from various disciplines to ensure the proper description of taxa, taxonomic stability over time, and preventing taxonomic inflation by integrating multiple lines of evidence [18, 20]. A “fast-track” taxonomy proposal arose from the pressing need to characterize and describe as much diversity as possible before it disappears, significantly shortening the description time by deliberately omitting previously collected material and reducing the most time-consuming parts of the process (i.e., high-quality images vs. detailed and extensive descriptions, reducing descriptions vs. enhance diagnoses) [21–22]. However, considering the overwhelming number of candidate species awaiting description worldwide (e.g., [23–31]), the global richness estimates [15], and the escalating biodiversity crisis, this strategy is likely unable to keep pace with current and projected extinction rates [32], nor to achieve a representative inventory of global species within a reasonable time frame [22]. Some authors, not without receiving criticisms, have attempted the so-called “turbo-taxonomy”, which consists in describing large numbers of taxa providing minimal information on each species, sometimes even just a barcoding sequence (e.g., [33]). While this approach may help overcoming the Linnean shortfall, questions arise about the usefulness and reliability of such species “descriptions” [34]. Although “fast-track” and “turbo” taxonomy are sometimes seen as synonyms [34], we distinguish them as two different approaches. We consider as “fast-track” those descriptions that, while concise, include all necessary sections to allow a third party to identify the species without the need for a molecular lab (e.g., [35]), while “turbo” are those minimalist descriptions that do not (e.g., [33]).

Regardless of the chosen approach, describing a species involves five steps, each depending on the previous one —the five D’s of taxonomy [36]: discovery, delimitation, diagnosis, description and specimen determination (although not necessarily by the same person). None of these steps can be achieved without previous field work. Field campaigns are time-consuming and, particularly in tropical regions, can be challenging, expensive, and logistically complex. To decide whether a lineage merits to be named, data from different lines of evidence should ideally be gathered, which requires time and cannot always be achieved within a single campaign. Compiling such evidence, preferably including multiple specimens for the type series, can be challenging when cryptic, secretive, seasonal, narrow-distributed species, or those with low population densities are involved. This is especially true for frogs, where obtaining call recordings

—often a source of key diagnostic characters— can be daunting. Specimens representing populations that clearly merit to be formally named can remain unstudied in shelves for years [37], which negatively impacts their conservation, as undescribed species are more vulnerable to extinction [38].

Amphibians are the most endangered vertebrate class worldwide. More than 41% of described species are under threat, and the number of species at high risk of extinction continues to increase [39]. Threats are ubiquitous throughout the amphibian tree of life, with habitat loss and degradation (primarily due to deforestation), climate change, and emerging diseases being the greatest risks to amphibian persistence [39]. There is a geographical bias, with those regions with greater diversity and lesser available information, i.e., the tropics, being the most affected [3, 40–41]. Paradoxically, current rates of amphibian species description are exponential and, during the last decade, hundreds of candidate species have been flagged globally [10, 30–31]. This indicates that we are still far from overcoming the Linnean amphibian shortfall, that the number of threatened species has likely been underestimated, and that many species are facing silent extinction before they are even discovered.

Here we analyze the type series metadata of all anuran species described since the year 2000 from four tropical regions, to globally and regionally estimate: (1) how long it takes to describe a frog since the first type specimen is collected using either traditional or contemporary methodologies?; (2) how long it takes to get the complete type series for describing a frog species?; (3) how long it takes to describe a frog species since all the type material has been collected?; (4) how many field campaigns are needed to describe a frog species?; (5) are the description times decreasing or increasing over time?; and (6) what abiotic factors determine that some frogs are described in a notorious shorter time than others, and how have these factors evolved over time?

## Materials and methods

This study did not involve any experiments or direct interaction with live animals. All data analyzed were obtained exclusively from previously published literature. Therefore, ethical approval from an Institutional Animal Care and Use Committee (IACUC) or other ethics board was not required for this research.

### Region selection, data acquisition and curation

We selected four regions across the globe based on two criteria: high frog species richness (i.e., > 300 species), and a current exponential rate of species description. The regions chosen were Ecuador, India, Madagascar, and Melanesia (Fig 1). These regions are taxonomically independent and, in principle, do not share taxonomists, which helps minimize regional biases in research practices. By selecting geographically scattered regions, we aim to investigate the time span of the species description process on a global scale, diluting regional biases to obtain a more accurate timeframe that can be considered globally applicable.

**Fig 1.**
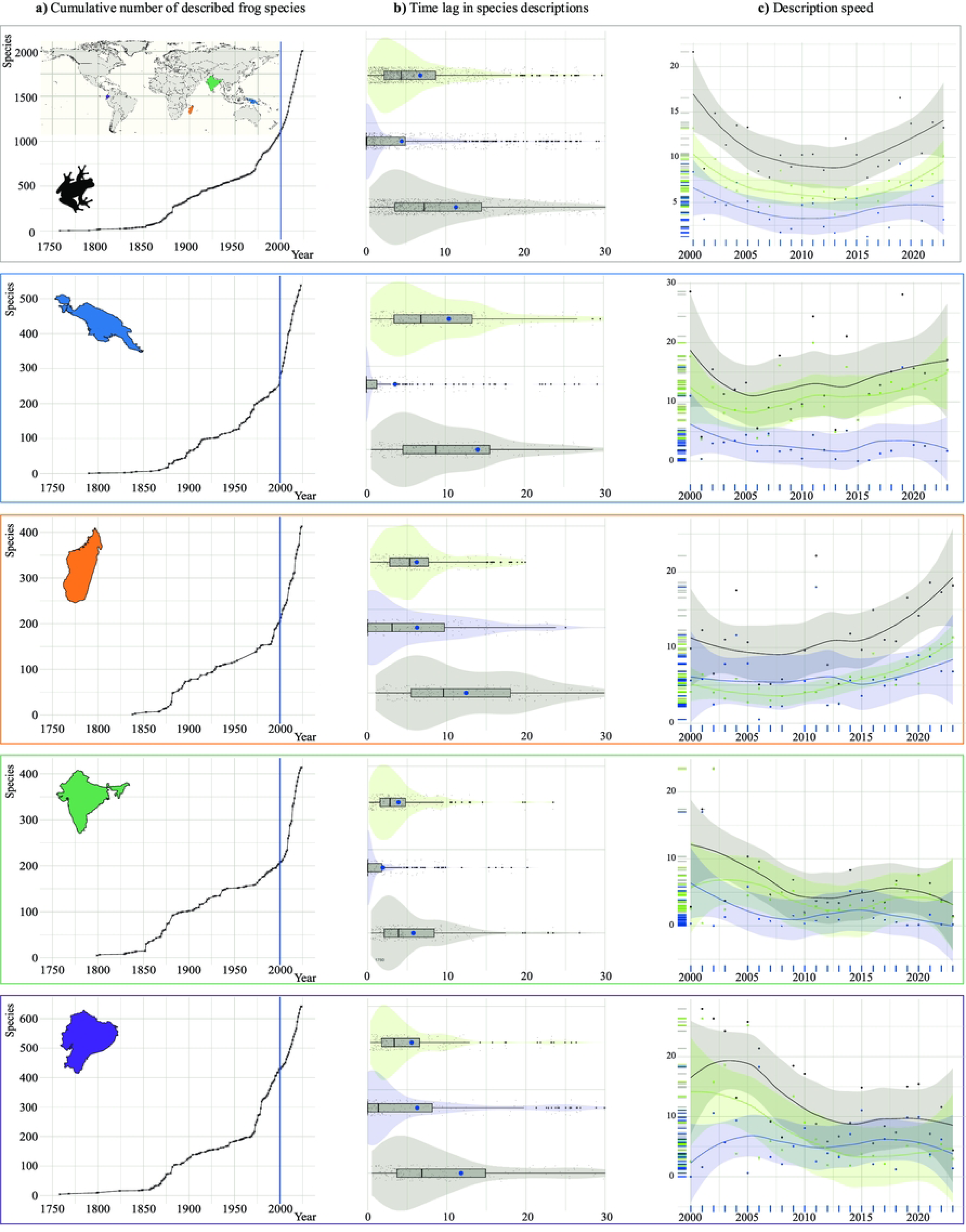
Time lags and trends in anuran amphibian species descriptions. (**a**) Cumulative number of frog species described over time. (**b**) Time lags in the description process. (**c**) Average description speed per year. Green: description *sensu stricto*, blue: collection, gray: overall process.

We retrieved the list of described frog species for each region, up to December 2023, from Amphibian Species of the World (ASW; [42]), filtering by higher taxa (Anura) and country, and retaining only native species (i.e., discarding the introduced). Since Melanesia includes two countries —Indonesia and Papua New Guinea—, and parts of Indonesia extend well beyond the region, we could not easily obtain the species list from the online database. Therefore, in accordance with the definition of the Melanesian region provided by Oliver et al. [31], we compiled the species list by selecting “Indonesia-Papua Region”, “Papua New Guinea”, “Solomon Islands”, and “Fiji” (which also includes smaller nearby islands) in ASW.

To determine the current species description lags (i.e., using contemporary methodologies), we filtered each regional species-list by year, retaining only those species described from the year 2000 (included) onward. We decided to exclude data predating 2000, as software and genetic data —assumed to accelerate species discovery, diagnosis, and description— have become widely available and utilized only in the 21st century. Genetic techniques have become crucial for describing species, but also for detecting cryptic species and screening regions (and public collections) to unveil overlooked and cryptic diversity (e.g., [10, 30]).

For each species, we retrieved the publication and all collection dates of the type series (holotype and paratypes) from the original descriptions, excluding additional and referred specimens. We only considered unique collection dates; that is, if multiple specimens were collected on the same day, we registered only one date. To deal with imprecise dates (e.g., 5-7 December 2008, February 2007) we used the midpoint date (e.g., 6 December 2008 and 14 February 2007, in the provided examples).

### Lag times

We made several necessary assumptions regarding the species description process, and defined three time periods: “overall description process”, “collection”, and “description *sensu stricto*”. We acknowledge that defining these periods and making these assumptions is an oversimplification, but it is necessary to draw conclusions (see Discussion).

We consider the “overall description process” as the time span between the collection date of the first (i.e., the oldest) specimen included in the type series and the publication date of the article introducing the new species to the scientific community. Within this overall process, we differentiate the two other sub-periods, the “collection” and the “description *sensu stricto*”.

The collection period refers to the field campaigns conducted to gather the entire type series, extending from the collection of the first specimen to the last. We assume that all type specimens were considered essential for the species description, which was delayed until the completion of the type series, and that the inclusion of both recently collected and older specimens was not arbitrary. Additionally, we regard the first collected type specimen as the first scientific encounter with the species (i.e., the moment a researcher truly discovers the species), even if it was not immediately recognized as a new taxon in the field and this insight occurred years later.

The description *sensu stricto* is the process encompassing taxon delimitation, diagnosis, description, and determination (*sensu* [36]), spanning from the collection of the last specimen of the type series to the publication date. At this point, we assumed that once the last specimen of the type series was collected, the description *sensu stricto* started.

To summarize and visualize the three periods, we calculated their durations in years for easier interpretation and representation. We obtained the central tendency and position statistics of each period using the *base* package in R [43], and we calculated asymmetry and dispersion statistics using the *psych* package [44]. We used the *ggplot2* package [45] to represent the results using violin and boxplots. This process was conducted independently for each genus and region, and merging the four regional datasets to obtain global estimates.

### Number of field campaigns and temporal patterns of type specimen collection

To estimate the number of field campaigns required to collect the entire type series for each species description, we calculated the time span between the sorted collection dates of the specimens in the type series. We considered as distinct field campaigns those collection events separated by more than one month (i.e., 30 days).

To estimate global and regional temporal patterns in the collection of type specimens (i.e., the number of field campaigns conducted to collect type specimens each month and year), we calculated the number of specimens collected per month and year, and represented the collection frequencies using heatmaps with the *ggplot2* package.

### Effect of abiotic variables on the description lags

To understand why certain species were described faster than others, we investigated the effect of several abiotic variables on the description *sensu stricto*. We did not analyze the effect of these variables on the overall and collection processes, as the values of these variables are typically determined post-collection.

We included three categorical variables: (1) whether genetics were involved in the species description (YES/NO), (2) the region, and (3) the genus; also, we considered five continuous variables: (1) the number of authors on the description paper, (2) the number of type specimens in the type series, (3) the number of species described in each paper, (4) the number of field campaigns required to collect the type series (see above), and (5) the number of species currently recognized in the genus.

Before fitting the models, we ensured that no numerical variable had a correlation greater than 70% with any other to avoid multicollinearity. For each of the four regions and the global analysis (i.e., combining the four regions into a single dataset), we fitted generalized linear mixed models (GLMM) using the *glmmTMB* R package [46]. In the global model, we included “genus” and “region” as random effects to account for the hierarchical structure of the data. We excluded “region” in the regional models. All models used a Gaussian family with an “identity” link function. We log-transformed the response variable to meet or improve the model assumptions.

To further assess the significance of the relationships, we performed an analysis of variance using the *Anova()* function from the *car* R package [47], which allowed us to evaluate the influence of the predictor variables without considering the factor levels.

We evaluated the model performance using the *DHARMa* package [48], by obtaining the simulated residuals from the model to assess the deviation from the expected distribution, uniformity, outliers, and dispersion. Additionally, we fitted null models (i.e., only including the random effects) and compared these with the empirical models to determine whether the inclusion of fixed effects in the empirical model, provided a significantly better fit to the data. We conducted this comparison using *anova()* from the *stats* package. Finally, we used the *effects* package [47] to represent the effects of the variables.

### Temporal evolution of variables

We evaluated the temporal evolution of several variables including the overall description process, the collection, the description *sensu stricto*, the number of authors, the number of species described each year, the number of taxonomic papers published annually, the number of species described in each paper, the number of type specimens in each species’ type series, the percentage of species described using genetics each year, and the number of field campaigns undertaken to collect the type series.

To analyze the temporal evolution of these variables, we averaged the yearly values for each region and globally (e.g., the mean time invested in the collection of type specimens for each species described in 2004); also, we counted the number of species described each year and the number of species described using genetics to compute the percentage.

We used the *ggplot2* package to generate all plots, applying local regressions (LOESS) to fit curves to the data for visualization purposes. To assess the temporal significance of the aforementioned variables, we calculated *p-values* using linear (LM, lm()) or generalized additive models (GAMs, gam()) from the *stats* package with default parameters [43]. We selected the best method based on the comparison of their coefficient of determination (R²).

## Results

### Species described since the year 2000 and lag times

Nine hundred and twenty-four frog species have been described across the four analyzed regions between 2000 and 2023. Melanesia leads with 286 newly described frog species, followed by Madagascar with 215, Ecuador with 213, and India with 210. This represents an average description rate of ca.10 new species per region each year. After excluding unavailable papers and species with unreported collection or publication dates, we analyzed a total of 2,981 unique collection dates from 896 species; 269 species from Melanesia, 215 from Madagascar, 208 from Ecuador, and 204 from India (S1 Table and S2 Table).

Thirty-six percent of the analyzed species were described within the first five years after the collection of the first type specimen, and ca. 26% of the species between five and ten years later.

The remaining ca. 40% of the species were described between ten and 125 years after the first type specimen collection (Fig 2).

**Fig 2.**
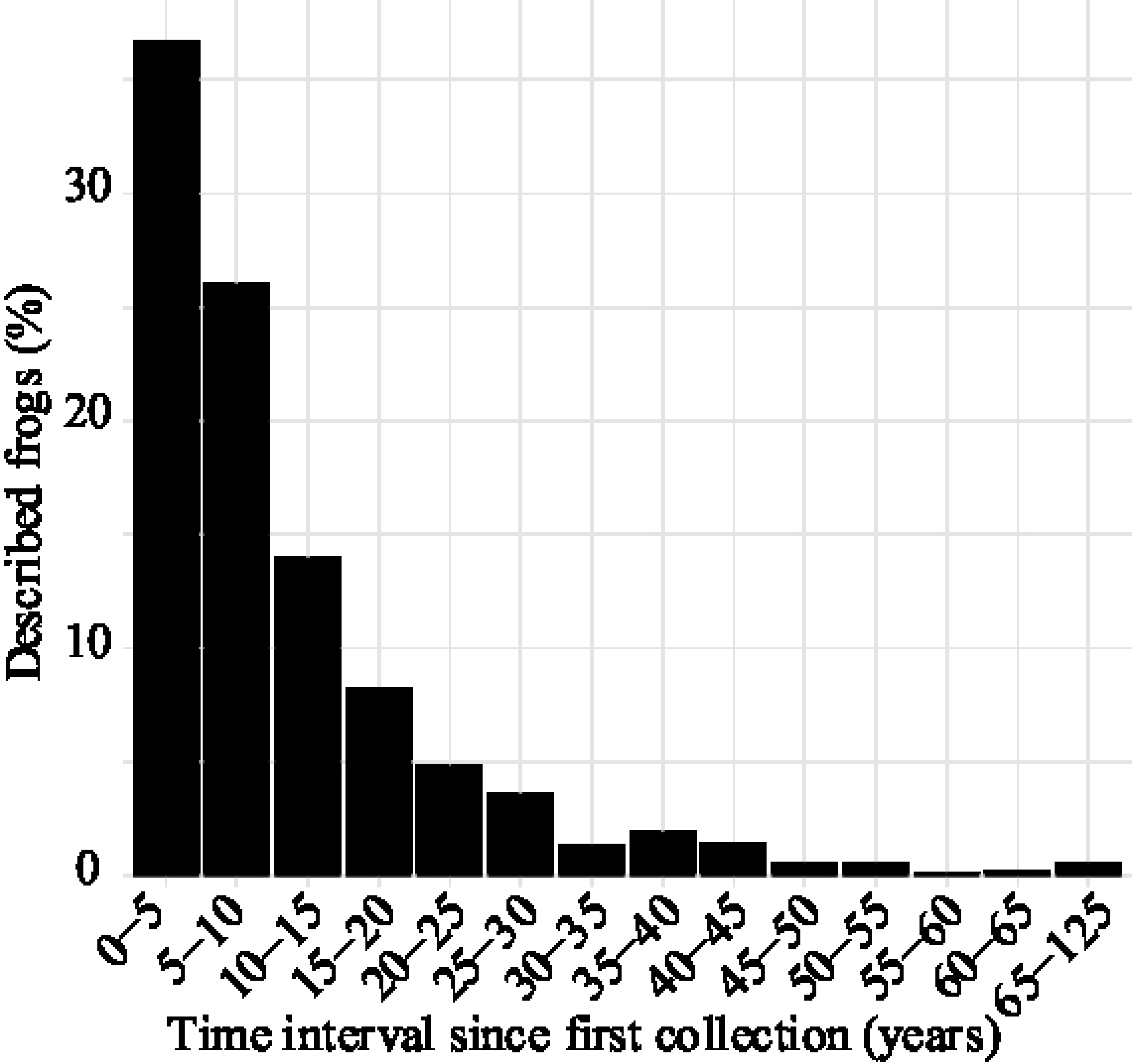
Percentage of frog species described since their first collection. Percentage of frog species described as a function of the time elapsed since the collection of the first specimen.

The mean and confidence interval for the global overall description process (i.e., from the collection of the first type-specimen to the paper publication date) is 11.26±0.82 years, with 4.47±0.65 years dedicated to collecting the type series and 6.79±0.49 years focused on the description *sensu stricto* (see Fig 1 and Table 1 for central tendency, position, asymmetry, and dispersion statistics).

**Table 1.** Summary statistics for each analyzed region across the different subperiods of the species description process. The table presents the minimum (Min.), mean, standard deviation (S.D.), 10% trimmed mean (Trim. mean), median, median absolute deviation (M.A.D), maximum (Max.), range, skewness (Skew), kurtosis, standard error (S.E.), and confidence interval for the mean (Interval).

|  | GLOBAL | Ecuador | India | Madagascar | Melanesia |
| --- | --- | --- | --- | --- | --- |
| Species | 896 | 208 | 204 | 215 | 269 |
| **Overall description process** |  |  |  |  |  |
| Min. | 0.42 | 0.5 | 0.42 | 0.99 | 0.61 |
| Mean | 11.26 | 11.79 | 5.77 | 12.43 | 14.08 |
| S.D. | 12.63 | 12.58 | 6.16 | 10.48 | 16.18 |
| Trim. mean | 8.91 | 9.43 | 4.78 | 11.25 | 11.01 |
| Median | 7.28 | 6.85 | 3.87 | 9.57 | 8.81 |
| M.A.D | 6.59 | 5.86 | 3.4 | 7.84 | 7.59 |
| Max | 125.74 | 82.95 | 60.04 | 110.86 | 125.74 |
| Range | 125.32 | 82.45 | 59.62 | 109.88 | 125.13 |
| Skew | 3.5 | 2.03 | 4.08 | 4.16 | 3.22 |
| Kurtosis | 20.48 | 5.17 | 28.75 | 34.34 | 15.31 |
| S.E. | 0.42 | 0.87 | 0.43 | 0.71 | 0.99 |
| Interval | $11.26 \pm 0.82$ | $11.79 \pm 1.71$ | $5.77 \pm 0.85$ | $12.43 \pm 1.40$ | $14.08 \pm 1.93$ |
| **Description *sensu stricto*** |  |  |  |  |  |
| Min | 0.18 | 0.33 | 0.18 | 0.35 | 0.57 |
| Mean | 6.79 | 5.54 | 3.88 | 6.19 | 10.45 |
| S.D. | 7.52 | 6.8 | 3.6 | 4.46 | 10.3 |
| Trim. mean | 5.33 | 4.15 | 3.27 | 5.56 | 8.43 |
| Median | 4.41 | 3.35 | 2.81 | 5.3 | 6.92 |
| M.A.D | 3.74 | 2.83 | 2.37 | 3.72 | 6.33 |
| Max | 54.62 | 51.48 | 23.44 | 19.91 | 54.62 |
| Range | 54.45 | 51.14 | 23.26 | 19.56 | 54.05 |
| Skew | 2.85 | 3.27 | 2.21 | 1.21 | 1.96 |
| Kurtosis | 10.57 | 14.09 | 6.8 | 0.82 | 3.93 |
| S.E. | 0.25 | 0.47 | 0.25 | 0.3 | 0.63 |
| Interval | 6.79 ± 0.49 | 5.54 ± 0.92 | 3.88 ± 0.49 | 6.19 ± 0.60 | 10.45 ± 1.23 |
| **Collection** |  |  |  |  |  |
| Min | 0 | 0 | 0 | 0 | 0 |
| Mean | 4.47 | 6.25 | 1.89 | 6.24 | 3.64 |
| S.D. | 9.93 | 10.58 | 5.13 | 9.96 | 11.54 |
| Trim. mean | 2.2 | 3.92 | 0.75 | 4.52 | 0.92 |
| Median | 0.08 | 1.32 | 0 | 3.07 | 0.01 |
| M.A.D | 0.12 | 1.96 | 0 | 4.56 | 0.02 |
| Max | 104.93 | 79.97 | 55.84 | 104.67 | 104.93 |
| Range | 104.93 | 79.97 | 55.84 | 104.67 | 104.93 |
| Skew | 4.94 | 2.84 | 6.54 | 5.05 | 5.51 |
| Kurtosis | 35.46 | 11.73 | 59.3 | 43.02 | 36.14 |
| S.E. | 0.33 | 0.73 | 0.36 | 0.68 | 0.7 |
| Interval | 4.47 ± 0.65 | 6.25 ± 1.44 | 1.89 ± 0.70 | 6.24 ± 1.33 | 3.64 ± 1.38 |

We found significant disparities among regions. India leads in the speed of species descriptions, averaging 5.77±0.85 years of overall process, 1.89±0.70 years for collection, and 3.88±0.49 years for description. Ecuador follows with an overall process time of 11.79±1.71 years, 6.25±1.44 years for collection, and 5.54±0.92 years for description. Madagascar closely trails, with an overall process of 12.43±1.4 years, comprising 6.24±1.33 years for collection and 6.19±0.60 years for description. Finally, the region that takes the longest is Melanesia, with an overall process time of 14.08±1.93 years, including 3.64±1.38 years for collection and 10.45±1.23 years for description (Fig 1 and Table 1).

In all three periods (i.e., overall, collection, and description) and across all five regions (i.e., four independent regions, plus global), we obtained high standard deviation values, with values exceeding the mean value in 11 out of 15 cases. This indicates a significant dispersion of the data around the mean. The median values consistently fell below the mean, suggesting a skewed distribution. Skewness values were greater than zero in all cases, pointing to positive asymmetry. Kurtosis values consistently exceeded zero, signaling heavy tails and the presence of more extreme values (outliers) that would typically occur in a normal distribution. Additionally, the standard error values remained low in all cases, especially in comparison to the standard deviation, indicating that the obtained mean is a fairly reliable estimate of the population mean.

### Number of field campaigns and temporal patterns of type specimen collection

Half of the species (476; 53.13%) described between 2000 and 2023 required a single field campaign to collect the entire type series, with an average of eight collected specimens per species. For 20.31% of species, two field campaigns were necessary, while 11.72% required three expeditions, and 6.14% required four. The rest (ca. 8%) of species needed between five and 23 field campaigns.

Considering the four regions altogether, species descriptions between 2000 and 2023 have been based on type specimens collected between 1893 and 2023. With over 60 type specimens collected each year, the years comprised between 1999 and 2016 show the highest concentration. Notably, February has been the peak month for type specimen collections, totaling 412 type specimens, followed by July with 277, November with 257, and June with 254 (S1 Fig).

In Madagascar, species descriptions have been based on specimens collected between 1900 and 2022, with a peak collection period from 2003 to 2007, during which more than 45 specimens were collected annually. February is, by far, the month with the highest number of type specimens collected, totaling 239 type specimens, followed by January with 125, November with 120, and March with 114 (S2 Fig).

Species descriptions in India have been based on specimens collected between 1954 and 2023, with the highest concentration between 2009 and 2012, with more than 25 specimens collected per year. June and July are the months with the highest number of type specimens collected, with a total of 82 type specimens collected in each month, followed by September with 53, August with 48, and May with 40 (S3 Fig).

In Melanesia, species descriptions have been based on specimens collected between 1893 and 2018, with the highest concentration between 2002 and 2005, with more than 65 specimens collected per year. April, May, September, and October are the months with the highest number of type specimens collected with between 104 and 109 specimens collected each month, followed by August with 85, July with 69, and November with 66 (S4 Fig).

Finally, the type specimens on which the species descriptions were based in Ecuador were regularly sampled between 1933 and 2022. The highest collection concentration is between 2000 and 2012, with more than 30 type specimens collected each year. July is the month with the highest number of type specimens collected with 110 type specimens, followed by February with 101, January with 86, and April with 82 (S5 Fig).

### Effect of abiotic variables on the description lags

Empiric models differed significantly from the null models and presented lower AIC values, indicating that the models are better than randomness and that the fixed variables have a genuine effect on the data (S3 Table). All models met the model assumptions, except for Ecuador, where minor violations were observed. Adding random effects (“genus” in all cases, and both “genus” and “region” in the global analysis) significantly increased the percentage of variance explained by the models (S4 Table, S6 Fig).

In all regions except Ecuador, the number of authors had a significant effect on species description time, with an increase in the number of authors associated with a faster description in the global model, Ecuador, and India, and with slower description times in Madagascar and Melanesia. The number of type specimens did not have a significant effect in any region. The number of described species within a genus was marginally significant in Madagascar, where richer genera are associated with faster species descriptions. This variable was not significant in other regions. The number of field campaigns significantly influences the description times in the global model and in India, was marginally significant in Ecuador and Madagascar, and showed no significant effect in Melanesia. Across all regions, however, an increase in the number of field campaigns to collect the type series was associated with shorter description times, perhaps because much of the work had already been done and only complementary data, such as vocalizations, were needed. The number of species described per paper was significant in the global model, Ecuador, and Melanesia, but not in Madagascar or India. An increase in the number of species per paper was associated with longer description times in all regions except in Madagascar, where the effect was the opposite (but not significant). The use of molecular data in species descriptions was significant in the global model and in India but not in Ecuador, Madagascar, or Melanesia. In all regions except Madagascar, incorporating genetic data was associated with longer description times (S7 Fig, S4 Table).

### Temporal evolution of variables

Globally, the overall description process decreased from 16 years in 2000 to 7.2 years by 2012. However, after 2012, the description lag increased again, reaching a current value of ca. 11 years. Both collection and description *sensu stricto* followed similar declining trends until 2012, after which the description time rose significantly, while collection time remained stable. Statistical analyses showed that both overall description time and description *sensu stricto* varied significantly with the year (p-value_GAM_: 0.023 and 0.015, respectively), while collection time remained stable (non-significant). In general, the overall description process has shortened since 2000, but recent trends suggest an ongoing increase (Fig 3).

**Fig 3.**
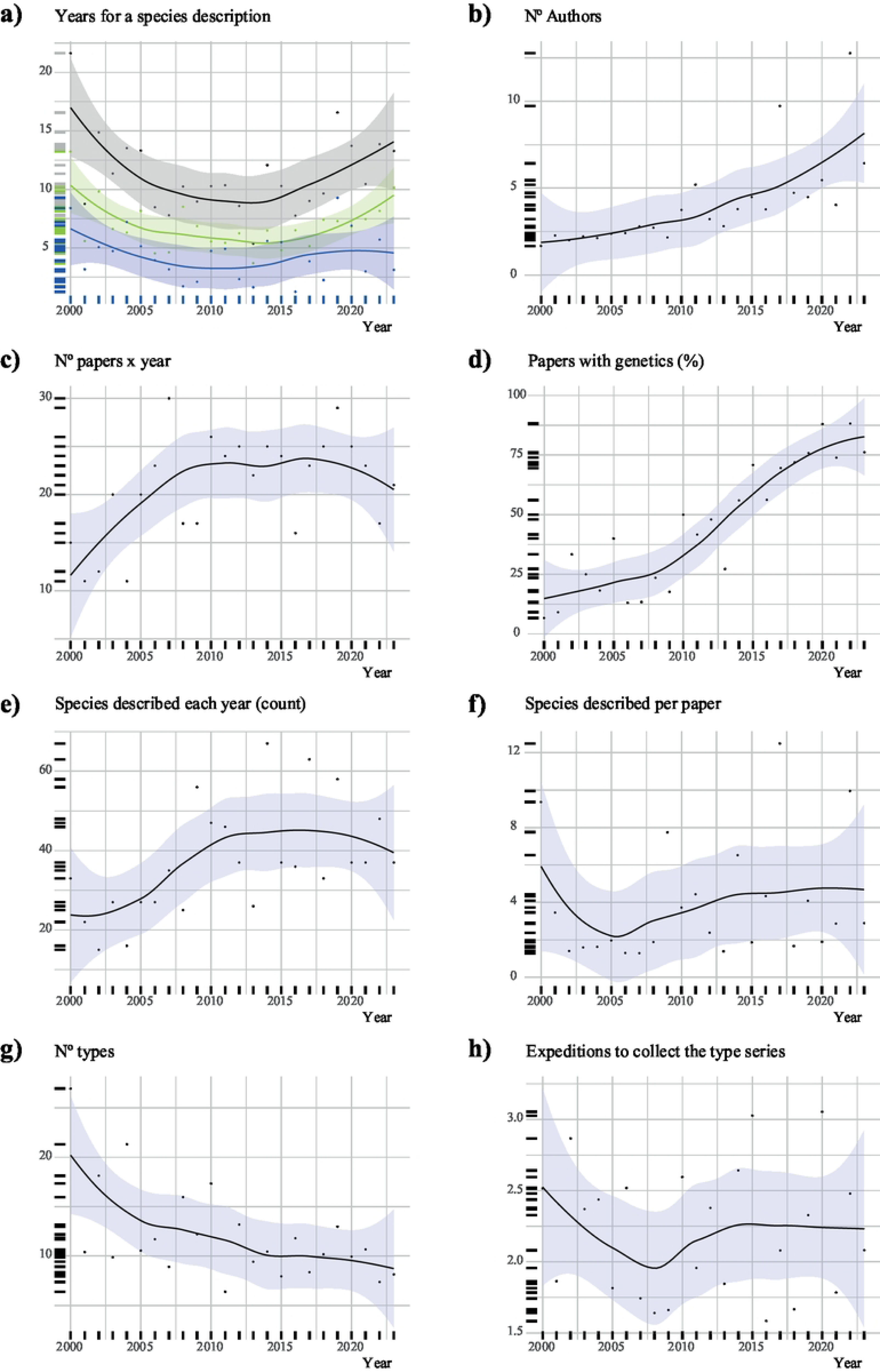
Temporal evolution of variables in the four analyzed regions. (**a**) Overall description process (black), collection (blue), and description *sensu stricto* (green), (**b**) N° of authors in each species description (mean), (**c**) Number of published taxonomic papers (count), (**d**) Taxonomic papers that include genetics (percentage), (**e**) N° of species described each year (count), (**f**) N° of species described in each paper (mean), (**g**) N° of type specimens in the type series (mean), (**h**) N° of field campaigns conducted to collect the entire type series (mean).

Ecuador and India have shown reductions in the overall description process between 2000 and 2012, after which they remained stable until recent declines (p-value_GAM_: 0.01 and 0.03, respectively). The description *sensu sticto* followed the same pattern as the overall description process in both regions (p-value_GAM_: 0.01 and non-significant, respectively). In India, the collection time has non-significantly decreased, while in Ecuador, it has remained stable. In contrast, Madagascar shows a considerable increase in both the overall description process and description *sensu stricto* (p-value_GAM_: 0.03 and <0.001, respectively) as well as a moderate non-significant increase in the collection time. Melanesia shows a similar pattern to global trends but non-significant, with an initial decrease followed by an increase in both the overall process and description *sensu stricto*, while the collection time has slightly declined (S8-S11 Fig).

The number of authors contributing to species descriptions has significantly increased globally (p-value_LM_: <0.001). In Ecuador and India, the number of authors rose from ca. two to ca. five authors, where it appears to have reached a plateau (p-values_GAM_: <0.001). In contrast, the number of authors in Madagascar and Melanesia continues to rise (p-values_GAM_: <0.006) (S8-S11 Fig).

The number of taxonomic papers published globally each year increased from ca. 12 to ca. 23, where it remained stable until 2020, when it began to decline again (p-value_GAM_: 0.01). Ecuador and India show similar patterns (p-values_GAM_: <0.001). In contrast, the number of papers in Madagascar shows high variability, with an overall mean of ca. five papers per year, and no significant trend. In Melanesia, there was also an increase followed by a decrease, but there is not a significant trend (S8-S11 Fig).

The proportion of taxonomic papers incorporating genetic data has significantly increased globally, reaching 95% of published papers using genetics in 2022 (p-value_GAM_: <0.001). Madagascar shows the earliest widespread usage of genetics, with 100% of published papers incorporating genetic data since 2004, with few exceptions (p-value_GAM_: <0.001). In India, the use of genetics remained low until 2010, when only 25% of papers incorporated genetic data. Since then, there has been a marked increase to the current 100% of usage (p-value_GAM_: <0.001). In Ecuador, the usage of genetics has increased steadily, currently exceeding 80% (p-value_GAM_: <0.001). In contrast, Melanesia lags behind the other regions, using genetics in only ca. 40% of the published papers, although there has been a modest increase (p-value: 0.005) (S8-S11 Fig).

The number of species described each year increased globally from ca. 25 to ca. 45 species until 2010, when it reached a plateau (p-value_GAM_: 0.01). A similar pattern is observed in Ecuador, where the plateau was reached in 2018, with ca. 12 species described annually (p-value_GAM_: <0.001). India shows a non-significant steady increase until 2012, followed by a decline. Madagascar and Melanesia show great variability with an overall non-significant continuous increase without reaching a plateau in Madagascar and an overall decrease in Melanesia (S8-S11 Fig).

The average number of species described per paper shows a globally non-significant positive relationship with the year. Ecuador and Madagascar exhibit similar patterns. In India and Melanesia, the number of described species per paper increased until 2012 and 2014, respectively, followed by a subsequent decline (S8-S11 Fig).

The number of type specimens in a type series has globally decreased from ca. 20 type specimens per species description to fewer than 10 (p-value_LM_ 0.005). A similar pattern is observed in Melanesia (p-value_LM_ 0.003). Ecuador and India do not show any significant trends in the evolution of the number type specimens. In Madagascar, the overall trend is a steady decrease, although some recently described species have significantly reversed that trend (p-value_GAM_ 0.004) (S8-S11 Fig).

The number of field campaigns required to collect the entire type series globally shows huge dispersion, with an overall stable trend between 1.5 and 3 field campaigns required to collect the type series (S8-S11 Fig).

## Discussion

### Anuran species descriptions amid a biodiversity crisis

Taxonomists have named more than 2.3 million species [49] at a pace of 20,000 species descriptions per year [50]. However, this represents only a small fraction of the world’s predicted species richness [15]. We are the first generation to fully comprehend that we are in the midst of an extinction event, while the majority of species remain undiscovered and undescribed. We may also be the last generation having the opportunity to explore and document Earth’s biodiversity [37, 51].

Here, we evaluated the time required to describe a frog species using either traditional and contemporary methodologies. Despite being the most endangered class of vertebrates [39], amphibians are being described at remarkably high rates, and the numerous candidate species flagged worldwide suggest that true amphibian richness —and consequently the number of threatened species— have been highly underestimated.

Our results suggest that the mean description time from the collection of the first type specimen to publication is 11.3 years per species, with 4.5 years dedicated to the type series collection and 6.8 years to the description *sensu stricto*. We found similar values across three of the four analyzed regions, with India showing lower values (see Results). Fontaine et al. [37] found similar “shelf” figures for tetrapods analyzing a random set of species described in 2007. Our results indicate that these figures are not decreasing at the global level. Our data suggest a trend of decreasing description times in continental regions (India and Ecuador), while description times are increasing in insular regions (Madagascar and Melanesia). This pattern may be explained by the greater presence and activity of local taxonomists in continental countries, while in the studied insular regions, taxonomy has historically been led by foreign institutions, which are experiencing a reduction in the number of specialists [52]. Overall, the global pattern is U-shaped, with current description times being longer than those in 2012 but shorter than those in 2000.

Only 36% of species described since 2000 were collected within a five-year timeframe, suggesting that new species are rarely identified in the field and that targeted sampling campaigns are uncommon. The remaining species were collected earlier —up to 125 years ago— and described much later. This underscores the importance of public collections as reservoirs of potential new species but, at the same time, it concerns about the lack of resources, time, or interest dedicated to examining the collected material. Public collections contain a wealth of overlooked specimens (e.g., [53–57]). In fact, as Bebber et al. [58] noticed, a significant percentage of the world’s undescribed species have already been collected and remain forgotten on museum shelves. In the current context of the biodiversity crisis, some authors have suggested prioritizing the collection and preservation of as many specimens as possible as testimonies of a vanishing biodiversity, ensuring future generations can describe species that may already be extinct [37, 59]. However, unless this strategy is paired with thorough revision of collected material, it may result in decades of specimens remaining unexamined on shelves, delaying species descriptions that could contribute to their conservation. To ensure species are described while they still exist, it is necessary to adopt streamlined species description methodologies (e.g., “fast-track taxonomy”), the integration of molecular data to support rapid identifications, and increasing funding for taxonomic research. Molecular screenings should become a standard practice, alongside proper processing of all collected specimens, at least to identify taxa that cannot be assigned to known species, especially after collection campaigns in poorly surveyed regions.

Earth’s tropical regions remain largely unexplored. Large-scale molecular screenings are still relatively rare, but those conducted have revealed high numbers (even hundreds) of potential new species [10, 30, 60]. Although the use of molecular data in species descriptions has increased in the last decades, it has yet to become a standard practice (see results, [50]). Our results suggest that incorporating molecular data into frog descriptions significantly increases description time in some regions. While adding any procedure to the species descriptions inherently increases the time required, the increase of time can also be attributed to cryptic species, which may remain undescribed on shelves for years until identified molecularly. Therefore, by leveraging new methodologies and increasingly affordable molecular tools, the widespread use of barcodes on collected material should become a common practice among taxonomists [61]. Genetically characterizing the collected material and making the sequences with specific localities publicly available would not only highlight candidate species, thereby promoting their formal description, but also provide valuable data for other disciplines. Subsequent analyses could help reduce the Wallacean shortfall (i.e., limited knowledge about species’ geographical distributions) by discovering new localities and improving our understanding of intraspecific diversity (e.g., macrogenetics; [62]).

Given the number of species still awaiting discovery and description, it is evident that we need more effective solutions for inventory and protect Earth’s amphibian fauna before it vanishes. Over an 11-year span, vast areas of currently suitable habitats are likely to disappear if current rates of habitat destruction continue [63–64].

### Types of species descriptions and future perspectives

A foundational task of taxonomy is comparing the variability of target specimens against a set of related species to assess whether the observed variability justifies considering them as a distinct unit [65]. Early species classifications were constrained by the morphological traits available at the time, leading to minimalist and often rudimentary descriptions due to the limited methodologies and understanding [21, 66]. Today, species descriptions are in general more complex, meticulous, and precise. However, significant disparities exist in their current structure and length. Traditional taxonomic practice follows a comprehensive approach to species descriptions, which, over time, incorporated new methodologies and lines of evidence within an integrative framework. This often led to descriptions that span tens of pages for a single species (e.g., [67]). In contrast, “fast-track taxonomy” accelerates species description by focusing on essential diagnostic traits, providing concise descriptions while maintaining diagnostic accuracy, and incorporating high-quality images to compensate for reduced textual detail. This results in shorter species accounts, ultimately allowing the description of hundreds of species in a single article (e.g., [35, 68]). Even more, within the framework of “turbo-taxonomy” [69], some authors describe hundreds of species based only on consensus barcode DNA sequence or molecular diagnostic characters, using these as the principal basis of species delimitation and diagnosis (e.g., [33, 70]). However, this approach has been widely criticized, as reliance on mitochondrial DNA (mtDNA) or barcodes alone does not provide sufficient evidence for robust species delimitation [71].

There is an undeniable urgency to describe as many species as possible before they disappear. However, speeding up the process of naming species should not compromise the quality of species hypothesis [22, 34]. Overcoming the Linnean shortfall while ignoring the Wallacean shortfall (and other shortfalls; see [72]) makes little sense if the ultimate goal is to conserve species. Identifying species without understanding where they occur or how they persist in their ecosystems limits the effectiveness of conservation efforts. However, must every shortfall be addressed in the initial process of description? Certain standards are clearly necessary to ensure taxonomic descriptions remains consistent, comparable, and facilitate the work of future taxonomists. Although new methods have emerged to disentangle taxonomic challenges derived from poor descriptions or poorly preserved specimens (e.g., [73]), we still face the consequences of brief historical descriptions, scattered (or lost) type material, and imprecise type localities, which deter anyone from attempting certain taxonomic decisions [21, 66]).

Taxonomy is becoming increasingly multidisciplinary, integrating approaches and techniques from disciplines such as genetics(omics), biogeography, bioacoustics, morphometry, osteology (CT scans). While these approaches may enhance accuracy and provide more information about species, they often require complex techniques and tools that can discourage non-specialists (but see iTaxotools; [74]). But how much complexity is necessary for a species description? Must every bone be described, or all biogeographical affinities revealed upfront? Another challenge in taxonomy is that much of the original biological materials used in species descriptions are scattered across museum collections worldwide, making it difficult and expensive to revisit and re-study. While genetic data are now centralized and easy to retrieve (i.e., GenBank, [75]; BOLD, [76]), other crucial data produced during species descriptions are either absent from accessible repositories, incomplete, or insufficiently standardized, limiting their reuse [50–51]. Even when available, these data may be inaccessible to many due to paywalls, such as those in Zootaxa, one of the most popular taxonomic journals. This hinders taxonomists in resource-limited regions from accessing free descriptions to describe vanishing species in hyperdiverse tropical countries.

A potential suggested solution to enhance both data accessibility and the speed of species descriptions, without compromising species hypothesis is “wikifying” taxonomy (cybertaxonomy). This approach integrates traditional taxonomic practices with modern digital platforms. If centralized, it would allow for the dynamic updating and sharing of species descriptions and data, enhancing accessibility and collaboration within the scientific community. It would promote concise, versioned species pages that can be continuously updated with new information (e.g., DNA sequences, traits, and distributional data) improving the efficiency of biodiversity research [21, 77–78]. This strategy would allow the mass description of species following a (to be defined) minimum standard to ensure reliable descriptions, with the possibility of later adding information to improve them ([21, 50]; and references therein). Consequently, it would not be necessary to delay naming species to provide the most complete description possible. Species could be included in the conservation scheme from the outset, and although the act of naming itself remains static, their descriptions and biological content could remain dynamic and be updated over time.

Species can be protected through direct conservation efforts by targeting specific species or indirectly by safeguarding their habitats. Under current legislation, only species bearing a scientific name are recognized for conservation. Undescribed species are not assessed by the IUCN and are excluded from biological inventories used to prioritize conservation areas. With the current figures of species descriptions, if we must wait until a species is formally named to protect it, we will lose the race against extinction for many of them. In some cases, taxonomists describe species whose existence has been known (to specialists) for decades, and when they finally name them, they realize that the locality where it was first collected has been completely destroyed (e.g., [79–80]). This highlights a critical gap in conservation: undescribed species constitute the majority of Earth’s biodiversity, but lack formal recognition and, consequently, legal protection, making them more vulnerable to extinction [38].

Therefore, the biggest challenge faced by undescribed species is that current conservation legislation is (described)species-centered. Since descriptions are slow and take increasingly longer, many species do not (and will not) have any form of protection unless they happen to occur within a protected area by chance. A potential solution is to use cybertaxonomy to include undescribed species in the conservation scheme from the outset, or alternatively, to enhance a region-centered conservation approach (i.e., indirect conservation). In the latter, we propose accounting for species that have been known for years but remain unnamed—many of which we have substantial data on (e.g., [35]). These species could be used to prioritize conservation areas based on species “richness”, aligning with global targets such as the 30x30 initiative, which aims to protect 30% of the planet’s land and oceans by 2030 [81]. This approach would prioritize long-term conservation efforts by considering a more realistic diversity inventory. To achieve this, we suggest publishing all available information on undescribed species, such as DNA barcodes and distribution data, to make it available for regional or global studies. We acknowledge that this approach, in some occasions, could inflate the species inventory of a given area. However, the worst outcome would be conserving divergent conspecific lineages, which, in any case, represent important genetic variability that should be preserved.

### Taxonomy crisis

Taxonomy is a foundational science. Almost all biological studies take place downstream ([36, 51]). Contradictorily, is an undervalued biological discipline [59]. Taxonomic papers receive few citations and are often published in low-impact journals [82–83] threatening the careers of early-career researchers with low citation indices [84–85]. Additionally, funding for taxonomy-focused projects is steadily declining [51, 77, 86–87]. To secure funding and publication in high-impact journals, these projects often need to be framed within cutting-edge methodologies. This can result in using a sledgehammer to crack a nut, ultimately extending description time spans.

Whether taxonomists are facing extinction is hotly debated [52, 59, 83, 88–90]. Some authors argue that the number of authors involved in species descriptions has increased exponentially [52, 91–93]. Others contend that, despite this author inflation, not all authors listed in taxonomic papers can be considered taxonomists [89,93]. On a global scale, there is a regional turnover, with historically leading taxonomic institutions losing taxonomists, while other regions are increasing their numbers [7, 52]. Our results confirm that the number of authors involved in frog species descriptions has been steadily increasing. However, we found a globally significant relationship between the increase in the number of authors and a reduction in the description time. These results contradict the presumed undesirable effects of author inflation hypothesis, indicating that task distribution among authors may lead to more efficient species descriptions. However, we did not find this relationship in all regions, suggesting that such improved efficiency may depend on the particularities of each research group.

Our results suggest that, since ca. 2010, the number of published frog taxonomical papers, the number of frog species described each year, and the number of species described per paper have remained stable. This contrasts with the growth in scientific productivity, partially driven by advancements in computational power, new and affordable sequencing technologies, and data analysis software [50, 73–74, 94–96]. Unchanged species diagnosis and description procedures have been flagged as the main cause [21, 50]. Moreover, the number of species described per taxonomist has generally declined. While some authors have attributed this trend to a lack of new species to describe [21, 88, 91, 93], the large number of candidate species flagged annually and current rates of species descriptions suggest otherwise. This decline may reflect a growing shortage of taxonomic specialists and an increased participation in taxonomy of researchers from other fields [89, 93, 97]. This trend may be further exacerbated by the fact that describing a new species, while inherently engaging for most biologists, it is little rewarded. As a result, despite not being taxonomists, some researchers venture into describing one or a few species.

Working in consortiums has been proposed as a strategy to increase the number of species described per article, thus improving the impact metrics of both articles and journals. While this strategy may increase impact metrics, an issue remains: who will be the first author? Is it more advantageous for early-career researchers to be the first author of a less-cited article or to be one among many contributors in a consortium article? Regarding description time, our results suggest that while an increase in the number of authors is associated with shorter description times at a global scale, a higher the number of species described per paper (which is expected to increase the impact metric) has the opposite effect, with more species leading to longer times required to describe each one.

### Study caveats

We had to define two subperiods in the species description process and make oversimplified assumptions to analyze data and draw general conclusions. First, the oldest collected specimen in the type series may not represent the species’ first scientific encounter, as earlier specimens might have been overlooked (deliberately or not) or stored elsewhere. Even if it was the first encounter, it might not have been recognized as new in the field. In some cases, collectors are not researchers, and the collected material is often not examined immediately after fieldwork. Second, a species can be described using only the holotype (or even part of it). Since not all specimens in the type series are essential for the species description, the process might not be delayed until their collection. Third, including ancient specimens in the type series of species description was not always strictly necessary, and their inclusion extended the calculated lag time between the first collection and publication. Fourth, the description *sensu stricto* rarely begins immediately after the last type specimen is collected. There are multiple factors that can delay the description of a species for decades. Lastly, we acknowledge that our figures may contain a level of error affecting all species in different degrees, which we did not account for: the time span between manuscript submission and publication. This interval varies depending on the journal and the length of the review process, which can sometimes be lengthy. For the purpose of this study, we have assumed that this interval is part of the species description.

## Acknowledgements

We want to acknowledge colleagues who provided fruitful discussions on this issue in the past years.

## Supporting information

**S1 Fig. Heatmap of global specimen collections.** Heatmap showing the distribution of specimen collection events in the four analyzed regions over time. The x-axis represents the months of the year, while the y-axis represents the years of collection. The color intensity indicates the number of specimens collected in each time period.

**S2 Fig. Heatmap of specimen collections in Madagascar.** Heatmap showing the distribution of specimen collection events in Madagascar over time. The x-axis represents the months of the year, while the y-axis represents the years of collection. The color intensity indicates the number of specimens collected in each time period.

**S3 Fig. Heatmap of specimen collections in India.** Heatmap showing the distribution of specimen collection events in India over time. The x-axis represents the months of the year, while the y-axis represents the years of collection. The color intensity indicates the number of specimens collected in each time period.

**S4 Fig. Heatmap of specimen collections in Melanesia.** Heatmap showing the distribution of specimen collection events in Melanesia over time. The x-axis represents the months of the year, while the y-axis represents the years of collection. The color intensity indicates the number of specimens collected in each time period.

**S5 Fig. Heatmap of specimen collections in Ecuador.** Heatmap showing the distribution of specimen collection events in Ecuador over time. The x-axis represents the months of the year, while the y-axis represents the years of collection. The color intensity indicates the number of specimens collected in each time period.

**S6 Fig. Residual diagnostics from the DHARMa R package for the fitted models.**

**S7 Fig. Significant effect plots of variables on description *sensu stricto*.** Colored boxes differentiate the regions. Gray= global, Purple = Ecuador, Orange= Madagascar, Green = India, and Blue = Melanesia.

**S8 Fig. Temporal evolution of variables in Ecuador**. (**a**) Overall description process (black), collection (blue), and description *sensu stricto* (green), (**b**) N° of authors in each species description (mean), (**c**) Number of published taxonomic papers (count), (**d**) Taxonomic papers that include genetics (percentage), (**e**) N° of species described each year (count), (**f**) N° of species described in each paper (mean), (**g**) N° of type specimens in the type series (mean), (**h**) N° of field campaigns conducted to collect the entire type series (mean).

**S9 Fig. Temporal evolution of variables in India**. (**a**) Overall description process (black), collection (blue), and description *sensu stricto* (green), (**b**) N° of authors in each species description (mean), (**c**) Number of published taxonomic papers (count), (**d**) Taxonomic papers that include genetics (percentage), (**e**) N° of species described each year (count), (**f**) N° of species described in each paper (mean), (**g**) N° of type specimens in the type series (mean), (**h**) N° of field campaigns conducted to collect the entire type series (mean).

**S10 Fig. Temporal evolution of variables in Madagascar**. (**a**) Overall description process (black), collection (blue), and description *sensu stricto* (green), (**b**) N° of authors in each species description (mean), (**c**) Number of published taxonomic papers (count), (**d**) Taxonomic papers that include genetics (percentage), (**e**) N° of species described each year (count), (**f**) N° of species described in each paper (mean), (**g**) N° of type specimens in the type series (mean), (**h**) N° of field campaigns conducted to collect the entire type series (mean).

**S11 Fig. Temporal evolution of variables in Melanesia**. (**a**) Overall description process (black), collection (blue), and description *sensu stricto* (green), (**b**) N° of authors in each species description (mean), (**c**) Number of published taxonomic papers (count), (**d**) Taxonomic papers that include genetics (percentage), (**e**) N° of species described each year (count), (**f**) N° of species described in each paper (mean), (**g**) N° of type specimens in the type series (mean), (**h**) N° of field campaigns conducted to collect the entire type series (mean).

**S1 Table. Collection and publication dates for each species and region extracted from the literature.**

**S2 Table. Lag times of collection (MEX_MIN_YEAR_COLPUB), description *sensu stricto* (MIN_YEAR_COLPUB) and overall process (MAX_YEAR_COLPUB) for each species and region. Categorical and numerical variables used in models are also included.**

**S3 Table. Statistical comparison of AIC values between empiric (fixed + random variables) and null (random variables only) models.**

**S4 Table. p-values from the GLMM models and evaluation metrics for global and regional analyses.**

## References

1. Lees AC, Pimm SL. Species, extinct before we know them?. Curr Biol. 2015; 25(5): R177–R180.

2. McCallum ML. Vertebrate biodiversity losses point to a sixth mass extinction. Biodiv Cons. 2015; 24(10): 2497–2519.

3. Ceballos G, Ehrlich PR, Dirzo R. Biological annihilation via the ongoing sixth mass extinction signaled by vertebrate population losses and declines. PNAS. 2017; 114(30):E6089–E6096.

4. Ceballos G, Ehrlich PR. The misunderstood sixth mass extinction. Science. 2018; 360(6393): 1080– 1081.

5. Young HS, McCauley DJ, Galetti M, Dirzo, R. Patterns, causes, and consequences of anthropocene defaunation. Annu Rev Ecol Evol Syst. 2016; 47: 333–358.

6. Tracewski Ł, Butchart SH, Di Marco M, Ficetola GF, Rondinini C, Symes A, et al. Toward quantification of the impact of 21st-century deforestation on the extinction risk of terrestrial vertebrates. Conserv Biol. 2016; 30(5): 1070–1079.

7. Prathapan KD, Rajan PD. Advancing taxonomy in the global south and completing the grand Linnaean enterprise. Megataxa. 2020; 1(1): 73–77.

8. Button S, Borzée A. A new multi-metric approach for quantifying global biodiscovery and conservation priorities reveals overlooked hotspots for amphibians. [Preprint]. arXiv:2308.08829. 2023 [cited 2024 Dec 12]. Available from: https://arxiv.org/abs/2308.08829.

9. Finn C, Grattarola F, Pincheira-Donoso D. More losers than winners: investigating Anthropocene defaunation through the diversity of population trends. Biol Rev. 2023; 98(5):1732–1748.

10. Carné A, Vieites DR. A race against extinction: The challenge to overcome the Linnean amphibian shortfall in tropical biodiversity hotspots. Divers Distrib. 2024; 30(12): 1–13.

11. Burriel-Carranza B, Tejero-Cicuéndez H, Carné A, Mochales-Riaño G, Talavera A, Al Saadi S, et al. Integrating genomics and biogeography to unravel the origin of a mountain biota: The case of a reptile endemicity hotspot in Arabia. Syst Biol. 2024; syae032.

12. Lomolino MV. Conservation biogeography. In: Lomolino MV, Heaney LR, editors. Frontiers of biogeography: New directions in the geography of nature. Sunderland (MA): Sinauer; 2004. p. 293.

13. Whittaker RJ, Araújo MB, Jepson P, Ladle RJ, Watson JE, Willis KJ. Conservation biogeography: assessment and prospect. Divers Distrib. 2005; 11(1): 3–23.

14. Mora C, Tittensor DP, Adl S, Simpson AG, Worm B. How many species are there on Earth and in the ocean? PLoS Biol. 2011; 9(8): e1001127.

15. Larsen BB, Miller EC, Rhodes MK, Wiens JJ. Inordinate fondness multiplied and redistributed: the number of species on earth and the new pie of life. Q Rev Biol. 2017; 92(3): 229–265.

16. Mace GM. The role of taxonomy in species conservation. Phil Trans R Soc B. 2004; 359(1444): 711– 719.

17. Nori J, Semhan R, Abdala CS, Rojas-Soto O. Filling Linnean shortfalls increases endemicity patterns: conservation and biogeographical implications for the extreme case of *Liolaemus* (Liolaemidae, Squamata) species. Zool J Linn Soc. 2022; 194(2): 592–600.

18. Padial JM, Miralles A, De la Riva I, Vences M. The integrative future of taxonomy. Front Zool. 2010; 7(1): 1–14.

19. Scheffers BR, Joppa LN, Pimm SL, Laurance WF. What we know and don’t know about Earth’s missing biodiversity. Trends Ecol Evol. 2012; 27: 501–510.

20. Padial JM, Castroviejo-Fisher S, Koehler J, Vila C, Chaparro JC, De la Riva I. Deciphering the products of evolution at the species level: the need for an integrative taxonomy. Zool Scr. 2009; 38(4): 431–447.

21. Riedel A, Sagata K, Suhardjono YR, Tänzler R, Balke M. Integrative taxonomy on the fast track-towards more sustainability in biodiversity research. Front Zool. 2013; 10: 1–9.

22. Vences M. The promise of next-generation taxonomy. Megataxa. 2020; 1(1): 35–38.

23. Meegaskumbura M, Bossuyt F, Pethiyagoda R, Manamendra-Arachchi K, Bahir M, Milinkovitch MC, Schneider CJ. Sri Lanka: an amphibian hot spot. Science. 2002; 298(5592): 379–379.

24. Fouquet A, Gilles A, Vences M, Marty C, Blanc M, Gemmell NJ. Underestimation of species richness in Neotropical frogs revealed by mtDNA analyses. PLoS ONE. 2007; 2(10):e1109.

25. Clare EL, Lim BK, Fenton MB, Hebert PD. Neotropical bats: estimating species diversity with DNA barcodes. PLoS ONE. 2011; 6(7): e22648.

26. Funk WC, Caminer M, Ron SR. High levels of cryptic species diversity uncovered in Amazonian frogs. Proc R Soc B. 2012; 279(1734): 1806–1814.

27. Milá B, Tavares ES, Munoz Saldana A, Karubian J, Smith TB, Baker AJ. A trans-Amazonian screening of mtDNA reveals deep intraspecific divergence in forest birds and suggests a vast underestimation of species diversity. PLoS ONE. 2012; 7(7): e40541.

28. Tänzler R, Sagata K, Surbakti S, Balke M, Riedel A. DNA barcoding for community ecology-how to tackle a hyperdiverse, mostly undescribed Melanesian fauna. PLoS ONE. 2012; 7(1): e28832.

29. Veijalainen A, Wahlberg N, Broad GR, Erwin TL, Longino JT, Sääksjärvi IE. Unprecedented ichneumonid parasitoid wasp diversity in tropical forests. Proc R Soc Ser B Biol. Sci. 2012; 279(1748): 4694–4698.

30. Vacher JP, Chave J, Ficetola FG, Sommeria-Klein G, Tao S, Thébaud C, et al. Large-scale DNA-based survey of frogs in Amazonia suggests a vast underestimation of species richness and endemism. J Biogeogr. 2020; 47(8): 1781–1791.

31. Oliver PM, Bower DS, McDonald PJ, Kraus F, Luedtke J, Neam K, et al. Melanesia holds the world’s most diverse and intact insular amphibian fauna. Comm Biol. 2022; 5(1): 1182.

32. Cowie RH, Bouchet P, Fontaine B. The Sixth Mass Extinction: fact, fiction or speculation?. Biol Rev. 2022; 97(2): 640–663.

33. Sharkey MJ, Janzen DH, Hallwachs W, Chapman EG, Smith MA, Dapkey T, et al. Minimalist revision and description of 403 new species in 11 subfamilies of Costa Rican braconid parasitoid wasps, including host records for 219 species. ZooKeys. 2021; 1013:1–665.

34. Fernandez-Triana JL. Turbo taxonomy approaches: lessons from the past and recommendations for the future based on the experience with Braconidae (Hymenoptera) parasitoid wasps. ZooKeys. 2022; 1087: 199–220.

35. Rakotoarison A, Scherz M, Glaw F, Köhler J, Andreone F, Franzen M, et al. Describing the smaller majority: integrative taxonomy reveals twenty-six new species of tiny microhylid frogs (genus *Stumpffia*) from Madagascar. Vertebr Zool. 2017; 67: 271–398.

36. Favret C. The 5 ‘D’s of Taxonomy: A User’s Guide. Q Rev Biol. 2024; 99(3): 131–156.

37. Fontaine B, Perrard A, Bouchet P. 21 years of shelf life between discovery and description of new species. Curr Biol. 2012; 22(22): R943–R944.

38. Liu J, Slik F, Zheng S, Lindenmayer DB. Undescribed species have higher extinction risk than known species. Conserv Lett. 2022; 15(3): e12876.

39. Luedtke JA, Chanson J, Neam K, Hobin L, Maciel AO, Catenazzi A, et al. Ongoing declines for the world’s amphibians in the face of emerging threats. Nature. 2023; 622(7982): 308–314.

40. Stuart SN, Chanson JS, Cox NA, Young BE, Rodrigues AS, Fischman DL, et al. Status and trends of amphibian declines and extinctions worldwide. Science. 2004; 306: 1783–1786.

41. Stuart SN, Hoffmann M, Chanson JS, Cox NA, Berridge RJ, Ramani PP, Young BE, editors. Threatened Amphibians of the World. Barcelona, Spain: Lynx Edicions; Gland, Switzerland: IUCN; Arlington, Virginia, USA: Conservation International; 2008.

42. Frost DR. Amphibian Species of the World: An Online Reference. Version 6.2 [Internet]. 2024 [cited 2024 Dec 12]. Available from: https://amphibiansoftheworld.amnh.org/index.php

43. R Core Team. R: A language and environment for statistical computing. R Foundation for Statistical Computing; 2024. Available from: https://www.R-project.org/.

44. Revelle W. psych: Procedures for Psychological, Psychometric, and Personality Research. R package version 2.4.6. Northwestern University, Evanston, Illinois; 2024. Available from: https://CRAN.R-project.org/package=psych.

45. Wickham H. ggplot2: Elegant Graphics for Data Analysis. Springer-Verlag New York; 2016.

46. Brooks ME, Kristensen K, van Benthem KJ, Magnusson A, Berg CW, Nielsen A, et al. glmmTMB balances speed and flexibility among packages for zero-inflated generalized linear mixed modeling. The R Journal. 2017; 9(2): 378–400.

47. Fox J, Weisberg S. An R companion to applied regression. 3rd ed. Thousan Oaks (CA):Sage; 2019.

48. Hartig F. DHARMa: Residual Diagnostics for Hierarchical (Multi-Level/Mixed) Regression Models [Internet]. R package version 0.4.7. 2024 [cited 2024 Dec 12]. Available from: https://CRAN.R-project.org/package=DHARMa.

49. Bánki O, Roskov Y, Döring M, Ower G, Hernández Robles DR, Plata Corredor CA, et al. Catalogue of Life [Internet]. Version 2024-07-18. Amsterdam (Netherlands): Catalogue of Life; 2024 [cited 2024 Dec 12]. Available from: 10.48580/dgbqz.

50. Miralles A, Bruy T, Wolcott K, Scherz MD, Begerow D, Beszteri B, et al. Repositories for taxonomic data: where we are and what is missing. Syst Biol. 2020; 69(6): 1231–1253.

51. Wheeler QD, Raven PH, Wilson EO. Taxonomy: impediment or expedient?. Science. 2004; 303(5656): 285–285.

52. Costello MJ, May RM, Stork NE. Can we name Earth’s species before they go extinct? Science. 2013; 339(6118): 413–416.

53. Sánchez-Vialas A, Calvo-Revuelta M, Castroviejo-Fisher S, De la Riva I. Synopsis of the amphibians of Equatorial Guinea based upon the authors’ field work and Spanish natural history collections. Proc Calif Acad Sci. 2020; 66:137–230.

54. Sánchez-Vialas A, Calvo-Revuelta M, De la Riva I. Synopsis of the reptiles of Equatorial Guinea. Zootaxa. 2022; 5202(1):1–197.

55. Sánchez-Vialas A, Copete-Mosquera LA, Calvo-Revuelta M. Contributions to the amphibians and reptiles of Myanmar: insights from the Leonardo Fea legacy housed at the Museo Nacional de Ciencias Naturales of Madrid. Zootaxa. 2024, 5457(1): 1–64.

56. Brown RM, Calvo-Revuelta M, Goyes Vallejos J, Sánchez-Vialas A, De la Riva I. On the second (or, rather, the first) specimen of the recently described *Calliophis salitan* (Squamata: Elapidae), with the first report of the species from Mindanao Island, southern Philippines. Herpetology notes. 2021; 14: 1027–1035.

57. López-Estrada EK, Sánchez-Vialas A, Manzanilla J, Piñango C, Ruiz JL, García-París M. An overview of the taxonomy and geographic distribution of Venezuelan *Epicauta* (Coleoptera: Meloidae). Ann Zool. 2022; 72(1): 9–47.

58. Bebber DP, Carine MA, Wood JR, Wortley AH, Harris DJ, Prance GT, et al. Herbaria are a major frontier for species discovery. PNAS. 2010; 107(51): 22169–22171.

59. Engel MS, Ceríaco LM, Daniel GM, Dellapé PM, Löbl I, Marinov M, et al. The taxonomic impediment: a shortage of taxonomists, not the lack of technical approaches. Zool J Linn Soc. 2021; 193(2): 381–387.

60. Jaramillo AF, De la Riva I, Guayasamin JM, Chaparro JC, Gagliardi Urrutia G, Gutiérrez R, et al. Vastly underestimated radiation of Amazonian salamanders (Plethodontidae: *Bolitoglossa*) and implications about plethodontid diversification. Mol Phylogenet Evol. 2020; 149: 1–23.

61. Padial JM, De la Riva I. Integrative taxonomists should use and produce DNA barcodes. Zootaxa. 2007; 1586: 67–68.

62. Blanchet S, Prunier JG, De Kort H. Time to go bigger: emerging patterns in macrogenetics. Trends Genet. 2017; 33(9): 579–580.

63. Vieilledent G, Grinand C, Rakotomalala FA, Ranaivosoa R, Rakotoarijaona JR, Allnutt TF, et al. Combining global tree cover loss data with historical national forest cover maps to look at six decades of deforestation and forest fragmentation in Madagascar. Biol Conserv. 2018; 222: 189–197.

64. Silva Junior CH, Pessôa AC, Carvalho NS, Reis JB, Anderson LO, Aragão LE. The Brazilian Amazon deforestation rate in 2020 is the greatest of the decade. Nat Ecol Evol. 2021; 5(2): 144–145.

65. Mayr E. Principles of systematic zoology. McGraw-Hill; 1969.

66. Godfray HCJ. Challenges for taxonomy. Nature. 2002; 417(6884): 17–19.

67. Scherz MD, Daza JD, Köhler J, Vences M, Glaw F. Off the scale: a new species of fish-scale gecko (Squamata: Gekkonidae: *Geckolepis*) with exceptionally large scales. PeerJ. 2017; 5: e2955.

68. Riedel A, Narakusumo RP. One hundred and three new species of *Trigonopterus* weevils from Sulawesi. ZooKeys. 2019; 828: 1–153.

69. Butcher BA, Smith MA, Sharkey MJ, Quicke DL. A turbo-taxonomic study of Thai *Aleiodes* (*Aleiodes*) and *Aleiodes* (*Arcaleiodes*) (Hymenoptera: Braconidae: Rogadinae) based largely on COI barcoded specimens, with rapid descriptions of 179 new species. Zootaxa. 2012; 3457(1): 1–232.

70. Meierotto S, Sharkey MJ, Janzen DH, Hallwachs W, Hebert PD, Chapman EG, et al. A revolutionary protocol to describe understudied hyperdiverse taxa and overcome the taxonomic impediment. Dtsch Entomol. 2019; 66(2): 119–145.

71. Reyes-Velasco J. A revision of recent taxonomic changes to the eyelash palm pitviper, *Bothriechis schlegelii* (Serpentes, Viperidae). Herpetozoa. 2024; 37: 305–318.

72. Hortal J, de Bello F, Diniz-Filho JAF, Lewinsohn TM, Lobo JM, Ladle RJ. Seven shortfalls that beset large-scale knowledge of biodiversity. Annu Rev Ecol Evol. Syst. 2015; 46(1): 523–549.

73. Rancilhac L, Bruy T, Scherz MD, Pereira EA, Preick M, Straube N, et al. Target-enriched DNA sequencing from historical type material enables a partial revision of the Madagascar giant stream frogs (genus *Mantidactylus*). J Nat Hist. 2020; 54(1–4): 87–118.

74. Vences M, Miralles A, Brouillet S, Ducasse J, Fedosov A, Kharchev V, et al. iTaxoTools 0.1: Kickstarting a specimen-based software toolkit for taxonomists. Megataxa. 2021; 6:77–92.

75. Benson DA, Karsch-Mizrachi I, Lipman DJ, Ostell J, Sayers EW. GenBank. Nucleic Acids Res. 2010; 38: 46–51.

76. Ratnasingham S, Hebert PD. BOLD: The Barcode of Life Data System (http://www.barcodinglife.org). Mol Ecol Notes. 2007; 7(3): 355–364.

77. Godfray Jr HCJ. Linnaeus in the information age. Nature. 2007; 446(7133): 259–260.

78. Wheeler QD, Knapp S, Stevenson DW, Stevenson J, Blum SD, Boom BM, et al. Mapping the biosphere: exploring species to understand the origin, organization and sustainability of biodiversity. Syst Biodivers. 2012; 10(1): 1–20.

79. Barratt CD, Lawson LP, Bittencourt-Silva GB, Doggart N, Morgan-Brown T, Nagel, P, et al. A new, narrowly distributed, and critically endangered species of spiny-throated reed frog (Anura: Hyperoliidae) from a highly threatened coastal forest reserve in Tanzania. Herpetol J. 2017; 27(1):13–24.

80. Vences M, Köhler J, Scherz MD, Hutter CR, Maheritafika HMR, Rafanoharana JM, et al. Four new species of forest-dwelling mantellid frogs from Madagascar allied to *Gephyromantis moseri*. Spixiana. 2024; 46(2):297–319.

81. Dinerstein E, Vynne C, Sala E, Joshi AR, Fernando S, Lovejoy TE, et al. A global deal for nature: Guiding principles, milestones, and targets. Sci Adv. 2019; 5(4): eaaw2869.

82. Agnarsson I, Kuntner M. Taxonomy in a changing world: seeking solutions for a science in crisis. Syst Biol. 2007; 56(3): 531–539.

83. Drew LW. Are We Losing the Science of Taxonomy?. BioScience. 2011; 61(12): 942–946.

84. Krell FT. Why impact factors don’t work for taxonomy. Nature. 2002; 415(6875): 957–957.

85. Walter DE, Winterton S. Keys and the crisis in taxonomy: extinction or reinvention?. Annu Rev Entomol. 2007; 52(1): 193–208.

86. Britz R, Hundsdörfer A, Fritz U. Funding, training, permits—the three big challenges of taxonomy. Megataxa. 2020; 1(1): 49–52.

87. Löbl I, Klausnitzer B, Hartmann M, Krell FT. The silent extinction of species and taxonomists—An appeal to science policymakers and legislators. Diversity. 2023; 15(10):1053.

88. Tancoigne E, Dubois A. Taxonomy: no decline, but inertia. Cladistics. 2013; 29(5): 567–570.

89. De Carvalho MR, Ebach MC, Williams DM, Nihei SS, Trefaut Rodrigues M, Grant T, et al. Does counting species count as taxonomy? On misrepresenting systematics, yet again. Cladistics. 2014; 30(3): 322–329.

90. Bradford-Grieve J. Is there a taxonomic crisis?. New Zealand Science Review. 2016; 73(3–4): 83–86.

91. Joppa LN, Roberts DL, Pimm SL. The population ecology and social behaviour of taxonomists. Trends Ecol Evol. 2011; 26(11): 551–553.

92. Bacher S. Still not enough taxonomists: reply to Joppa et al. Trends Ecol Evol. 2012; 27(2): 65–66.

93. Bebber DP, Wood JR, Barker C, Scotland RW. Author inflation masks global capacity for species discovery in flowering plants. New Phytol. 2014; 201(2):700–706.

94. W etterstrand K. DNA sequencing costs [Internet]. Available from: www.genome.gov/sequencingcostsdata. [cited 2024 Nov 26].

95. Puillandre N, Brouillet S, Achaz G. ASAP: assemble species by automatic partitioning. Mol Ecol Resour. 2021; 21(2): 609–620.

96. Ramírez JL, Valdivia P, Rosas-Puchuri U, Valdivia NL. SPdel: A pipeline to compare and visualize species delimitation methods for single-locus datasets. Mol Ecol Resour. 2023; 23(8): 1959–1965.

97. Tancoigne E, Ollivier G. Evaluating the progress and needs of taxonomy since the Convention on Biological Diversity: going beyond the rate of species description. Aust Syst Bot. 2017; 30(4): 326– 336.

